# High-resolution double vision of the archetypal protein tyrosine phosphatase

**DOI:** 10.1101/2023.09.12.557361

**Authors:** Shivani Sharma, Tamar (Skaist) Mehlman, Reddy Sudheer Sagabala, Benoit Boivin, Daniel A. Keedy

## Abstract

Protein tyrosine phosphatase 1B (PTP1B) plays important roles in cellular homeostasis and is a highly validated therapeutic target for multiple human ailments including diabetes, obesity, and breast cancer. However, much remains to be learned about how conformational changes may convey information through the structure of PTP1B to enable allosteric regulation by ligands or functional responses to mutations. High-resolution X-ray crystallography can offer unique windows into protein conformational ensembles, but comparison of even high-resolution structures is often complicated by differences between datasets including non-isomorphism. Here we present the highest-resolution crystal structure of apo wildtype (WT) PTP1B to date, out of ∼350 total PTP1B structures in the PDB. Our structure is in a crystal form that is rare for PTP1B, with two unique copies of the protein that exhibit distinct patterns of conformational heterogeneity, allowing a controlled comparison of local disorder across the two chains within the same asymmetric unit. We interrogate the conformational differences between these chains in our apo structure, and between several recently reported high-resolution ligand-bound structures. We also examine electron density maps in a high-resolution structure of a recently reported activating double mutant, and discover unmodeled alternate conformations in the mutant structure that coincide with regions of enhanced conformational heterogeneity in our new WT structure. Our results validate the notion that these mutations operate by enhancing local dynamics, and suggest a latent susceptibility to such changes in the WT enzyme. Together, our new data and analysis provide a freshly detailed view of the conformational ensemble of PTP1B, and highlight the utility of high-resolution crystallography for elucidating conformational heterogeneity with potential relevance for function.

## Introduction

Proteins are dynamic molecules that undergo continuing motion. While it offers no insights into the timescales of such motions, X-ray crystallography can reveal detailed, atomistic information about a protein’s conformational ensemble. Such information can be represented in the form of multiconformer models with local alternate conformations (Keedy, Fraser, and van den Bedem 2015; Riley et al. 2021; Wankowicz et al. 2023). The scale of the shifts between such alternate conformations can vary, ranging from small-scale backbone changes coupled to side-chain rotamer changes (Lovell et al.2000; Davis et al. 2006) to larger-scale backbone shifts of secondary structure or loops (Deis et al. 2014; Keedy et al. 2018), although it is worth noting that even small-scale changes in protein conformation can be critical for biological function (Barstow et al. 2008; Fraser et al. 2009; Yabukarski et al. 2020). Moreover, the local conformations of neighboring residues in proteins depend upon one another (Martin et al. 2011; van den Bedem et al. 2013; Bhattacharyya, Ghosh, and Vishveshwara 2016; Johansson and Lindorff-Larsen 2018). Importantly, even apo (unliganded) proteins are prone to sample low-occupancy conformations that change in population and contribute to function in response to molecular events (Keedy et al. 2018; Wankowicz et al. 2022; Greisman, Dalton, et al. 2023).

Despite the allure of X-ray crystallography for deciphering conformational heterogeneity in proteins, it has some technical limitations. First, resolution can be suboptimal, limiting our ability to resolve low-occupancy alternate conformations. Second, comparing two crystal structures from different experiments can be complicated by factors such as non-isomorphism, differences in crystallization conditions, cryocooling stochasticity (Keedy et al. 2014; Fischer, Shoichet, and Fraser 2015), and coordinate error for small changes (Davis et al. 2006). An ideal scenario would be a single crystal that diffracts to high resolution and reveals distinct protein states, thus allowing a controlled comparison between conformations and exploration of coupling between protein sites.

Here we present a new crystal structure of the dynamic allosteric enzyme PTP1B (PTPN1) (Keedy et al. 2018; Whittier, Hengge, and Loria 2013; Choy et al. 2017), the archetypal protein tyrosine phosphatase (PTP). Our structure is the highest-resolution (1.43 Å) structure of apo wildtype (WT) PTP1B to date, out of ∼350 structures of PTP1B deposited in the Protein Data Bank (PDB) (Berman et al. 2000). Furthermore, it assumes a rare crystal form displaying non-crystallographic symmetry with two distinct copies of the protein in different environments within the same crystal lattice. This crystal form had never been observed for this protein until very recently for a series of structures bound to small-molecule fragments (Greisman, Willmore, et al. 2023; Morris et al. 2023) — but has not yet been reported for the apo enzyme, and the conformational differences between the two distinct chains have not been studied. Another report included several structures of PTP1B with active-site mutations in the same space group, but they have different cell dimensions, only one copy per asymmetric unit, and lower resolution (Morris et al. 2023) so are not directly relevant here. We exploit the fortuitous arrangement within our crystal to directly compare distinctly ordered states of PTP1B in atomic detail, including conformational differences that span distal regions of the structure such as key allosteric sites (Keedy et al. 2018; Skaist Mehlman et al. 2023).

We also compare our new structure to other notable, recently published structures of PTP1B. This includes the recently published structures of WT PTP1B bound to small-molecule fragments at non-orthosteric sites with the same rare crystal form (Greisman, Willmore, et al. 2023). Those liganded structures were not accompanied by an apo structure, which we now provide. Our analysis also includes mutant structures of PTP1B, identified based on coevolution and designed to modulate dynamics (Torgeson et al. 2022). These structures include the only structure of PTP1B at higher resolution (1.24 Å) than ours (1.43 Å), but they do not include a structure of WT PTP1B, and do not exhibit non-crystallographic symmetry. Our reanalysis of these published structures has unearthed additional “hidden” conformational heterogeneity (Lang et al. 2010) that was previously left unmodeled and helps explain the functional effects of the mutations.

Overall, using our new high-resolution apo WT structure, we observe variable levels of conformational disorder in one protein chain versus another, with the effect notably more pronounced for allosteric regions. We also highlight instances of coupled alternate conformations, wherein one residue becomes more flexible while a neighboring residue becomes less flexible. Finally, we report a striking colocalization of (i) coupled alternate conformations in the apo state, (ii) activating mutations with surrounding residues exhibiting previously unmodeled structural responses, and (iii) nearby small-molecule fragment binding. Together, our new data and reanalysis of other recent data hint at an even broader allosteric network in PTP1B than previously realized (Keedy et al. 2018; Choy et al.2017), and highlight the value of high-resolution crystallography and multiconformer modeling for obtaining unique windows into protein conformational ensembles that may pertain to function.

## Results

### Unique crystallographic dataset for PTP1B

We have determined a new crystal structure of apo WT PTP1B to high resolution (1.43 Å) (**Table 1**). Our structure is the highest-resolution apo structure of WT PTP1B to date; the second-highest resolution for a deposited apo structure of WT PTP1B is 1.50 Å (PDB ID: 2cm2) (Ala et al. 2006). Moreover, the space group of our dataset is P 43 21 2, which is rare for PTP1B. Indeed, among the ∼350 crystal structures of PTP1B deposited in the PDB, spanning seven different space groups, only five total deposited structures (excluding ours) have the same crystal form (with the same cell dimensions) as our structure, albeit all with bound ligands and from one recent study (Greisman, Willmore, et al. 2023). Thus, the structure we report here is the first of apo PTP1B in this new crystal form. Notably, this crystal form includes two non-identical chains in the asymmetric unit. In the sections to follow, we interrogate differences in conformational heterogeneity between these two chains in detail, with an eye toward potential involvement in allosteric mechanisms.

**Table 1:** Crystallographic statistics. Overall statistics given first (statistics for highest-resolution bin in parentheses).

|  |  |
| --- | --- |
| PDB ID | 8u1e |
| Resolution (Å) | 30.59–1.43 (1.48–1.43) |
| Completeness (%) | 97.94 (85.28) |
| Multiplicity | 25.6 (25.7) |
| I/sigma(I) | 10.82 (0.31) |
| R<sub>merge</sub>(I) | 0.147 (9.777) |
| R<sub>meas</sub>(I) | 0.150 (9.974) |
| R<sub>pim</sub>(I) | 0.029 (1.955) |
| CC<sub>1/2</sub> | 1.000 (0.404) |
| Wilson B-factor (Å<sup>2</sup>) | 24.84 |
| Total observations | 3,052,767 (302,442) |
| Unique observations | 119,377 (11,763) |
| Space group | P 43 21 2 |
| Unit cell dimensions<br>(Å, Å, Å, °, °, °) | 88.41, 88.41, 163.00,<br>90, 90, 90 |
| Solvent content (%) | 44.80 |
| R<sub>work</sub> | 0.150 (0.352) |
| R<sub>free</sub> | 0.202 (0.392) |
| RMS bonds (Å) | 0.010 |
| RMS angles (°) | 1.04 |
| Ramachandran outliers (%) | 0.36 |
| Ramachandran favored (%) | 97.33 |
| Clashscore | 3.35 |
| MolProbity score | 1.25 |

### Global structural differences between chains

Numerous studies using X-ray crystallography data have established the role of crystal packing/contacts on the observed conformational landscape of a protein (Jacobson et al. 2002; Bhabha, Biel, and Fraser 2015; Tyka et al. 2011). Our new structure of PTP1B contains two nonidentical chains in the asymmetric unit (ASU), each experiencing distinct packing within the crystal lattice. In particular, chain A has fewer crystal contacts and only ∼40% as much total crystal contact surface area relative to chain B (**Table 2**). Consistent with the idea that more extensive crystal packing can stabilize protein conformations whereas less extensive packing can allow conformational disorder, chain A has higher average B-factors than chain B, whether considering Cα backbone atoms or all atoms including side chains (**Table 2**). These changes in protein disorder appear to be coupled to significant changes in solvation at the protein surface: strikingly, chain A has only ∼67% as many ordered water molecules as chain B (**Table 2**). Together, these observations illustrate how the two distinct chains in our new high-resolution structure provide a useful avenue to explore how local conformational changes — in this case from crystal contacts — can affect other parts of a protein structure.

**Table 2:** Differences in structural metrics between chains. See Methods for calculation details.

|  | Chain A | Chain B |
| --- | --- | --- |
| # of crystal contacts | 3 | 5 |
| Total crystal contact surface area ($\text{\AA}^2$) | 749.7 | 1869.7 |
| Average B-factor, Cα atoms ($\text{\AA}^2$) | 35.5 | 26.7 |
| Average B-factor, all atoms ($\text{\AA}^2$) | 38.6 | 29.5 |
| # of ordered water molecules | 156 | 233 |

The next part of our analysis focuses on specific regions with backbone movements between the two chains, based on the Cα-Cα distance after superposition. Interestingly, we observe that the residues with the largest differences across the two chains generally correspond to regions previously annotated as being allosteric, including Loop 16 of the L16 allosteric site, the N-terminus adjacent to the L16 site, and the C-terminus including the allosteric α7 helix (**Fig. 1**). Several active-site loops including the WPD loop maintain a relatively similar conformation between chains.

**Figure 1:**
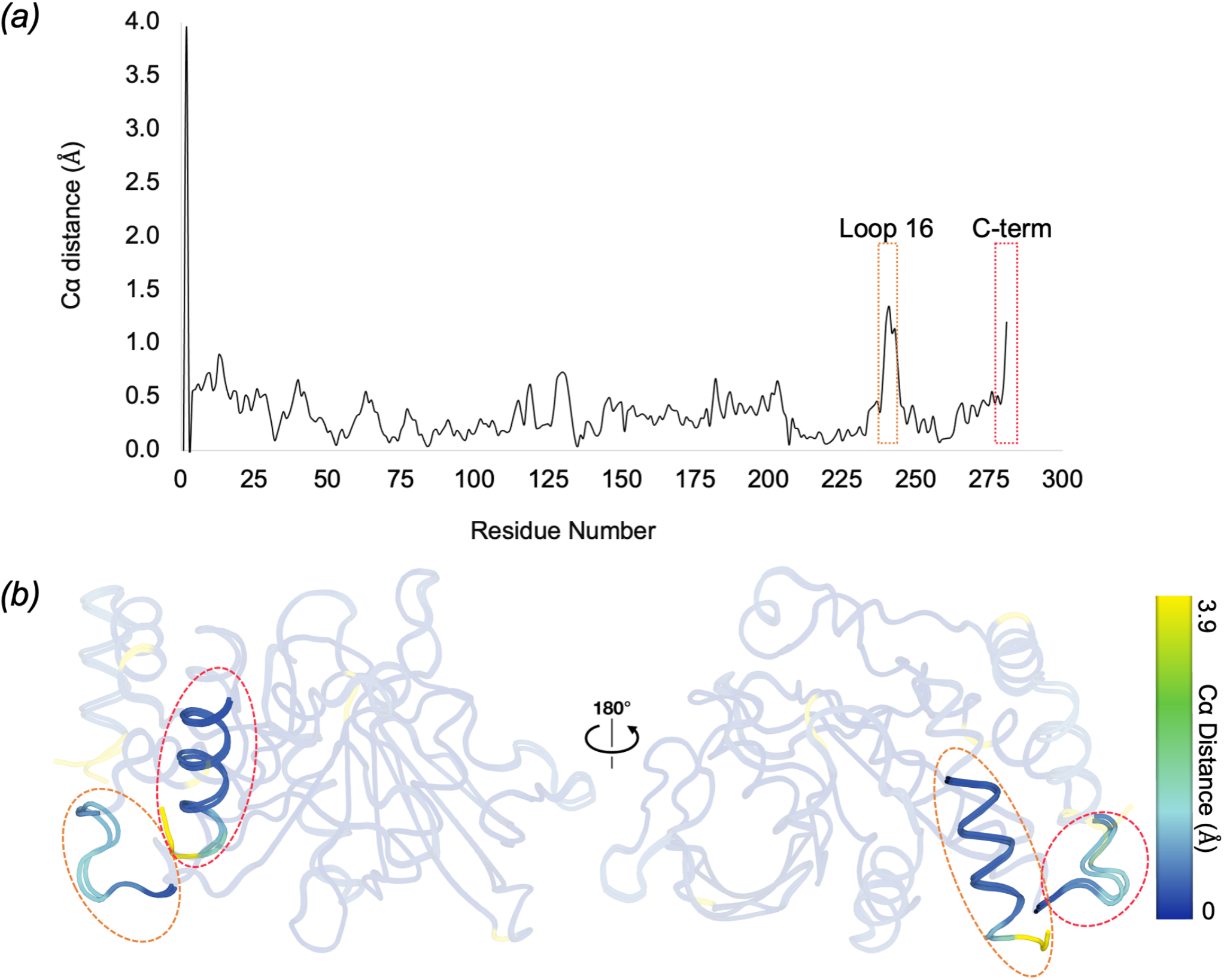
Sites displaying high backbone displacement between two distinct protein chains. *(a)*Plot of inter-chain Cα distance vs. amino acid sequence.*(b)*Overlay of both chains with residues colored by inter-chain Cα distance, viewed from two different angles. Regions of interest in *(a)* and *(b)* are highlighted with colored dashed outlines.

We also performed the same analysis with the only other five structures of PTP1B in the same crystal form (all ligand-bound) (PDB IDs: 8g6a, 8g65, 8g67, 8g68, 8g69) (Greisman, Willmore, et al. 2023). The results point to the same allosteric regions as with our new structure, albeit with even greater effects for some structures in some regions such as the L16 site (**Fig. S1**). The results for the ligand-bound structures also point to some additional areas including the active-site E loop (residues ∼110–120) and the region near residues 60–65, both of which can be difficult to model into local density and exhibit coordinate variability across various PTP1B structures in the PDB.

### Differences in disorder from B-factors

To test whether overall differences in disorder between the two chains (**Table 2**) are distributed evenly vs. heterogeneously throughout the protein structure, we studied B-factors on a per-residue basis. As the two chains are from the same crystal structure, no extra normalization of B-factors is required. A side-by-side comparison of the B-factors in the two chains of our structure implicates the active-site WPD loop, active-site E loop, and allosteric L16 site as exhibiting relatively high disorder (**Fig. 2 *(a)***). B-factor differences between chains indicate differential conformational effects in these regions: for example, the WPD loop is more flexible in chain A, whereas the L16 site is more flexible in chain B (**Fig. 2 *(b)***). Thus, despite chain A being more flexible overall, different local regions are more or less flexible in either chain.

**Figure 2:**
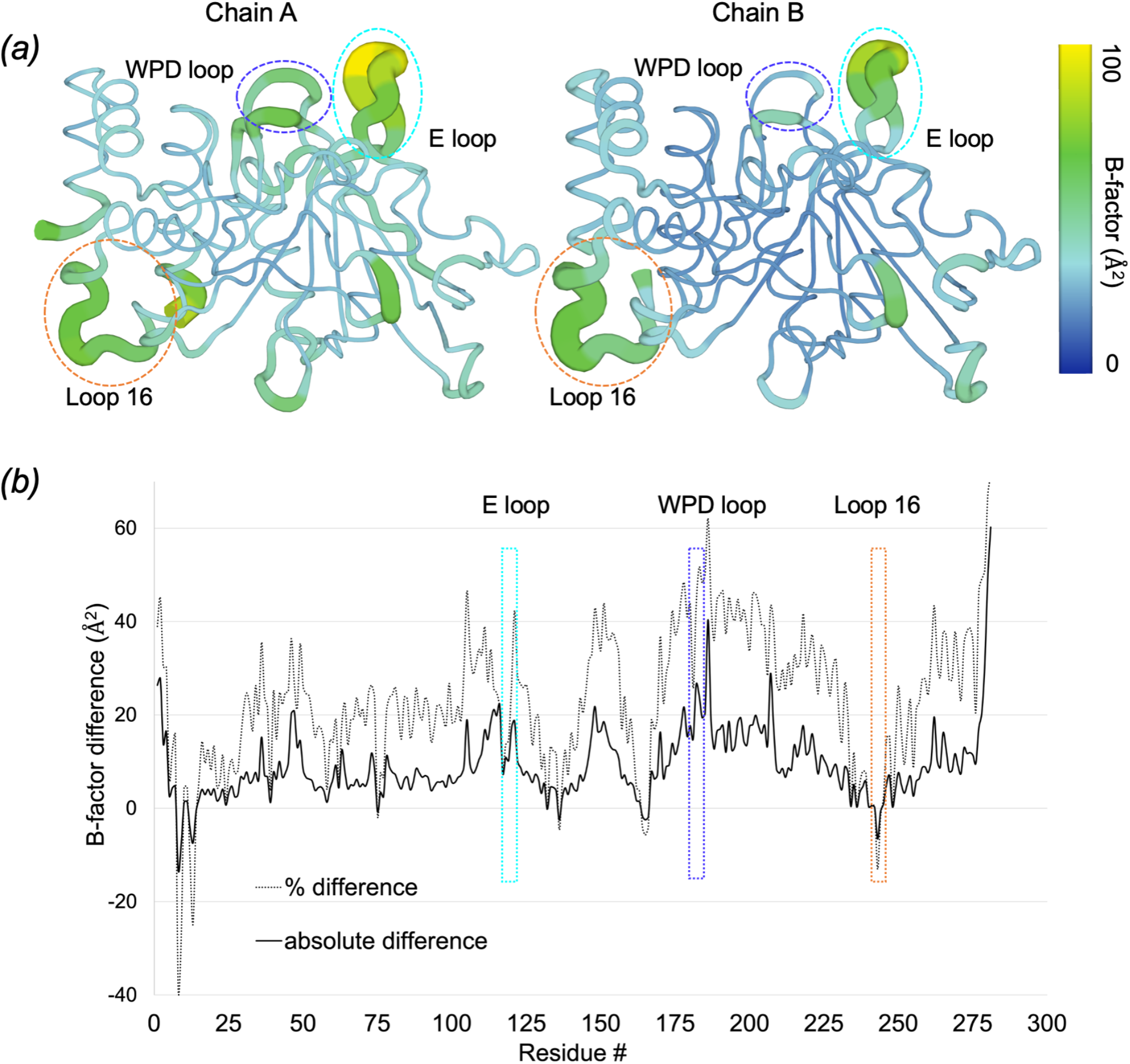
Differences in atomic disorder across sites and between distinct protein chains. *(a)*All-atom B-factors in chain A (left) vs. chain B (right). *(b)*All-atom absolute (solid line) and relative (%, dotted line) B-factor difference (chain A minus chain B) plotted as a function of amino acid sequence. In both panels, notable regions with high disorder and/or differential disorder between the two chains are highlighted with dashed lines (blue: WPD loop, cyan: E loop, pink: L16 allosteric site).

### Local structural differences between chains

A superposition of the two chains in our new structure reveals several differences in local conformations (**Fig. 3 *(a)***). First, although the WPD loop is in the open conformation in both chains of our structure, only in chain B does F182 (the F of the WPDFG motif (Yeh et al. 2023)) adopt a side-chain rotamer conformation that is quite rare for PTP1B (χ1 *t* near 180°) (Lovell et al. 2000) (**Fig. 3 *(b)*, middle panel**).

**Figure 3:**
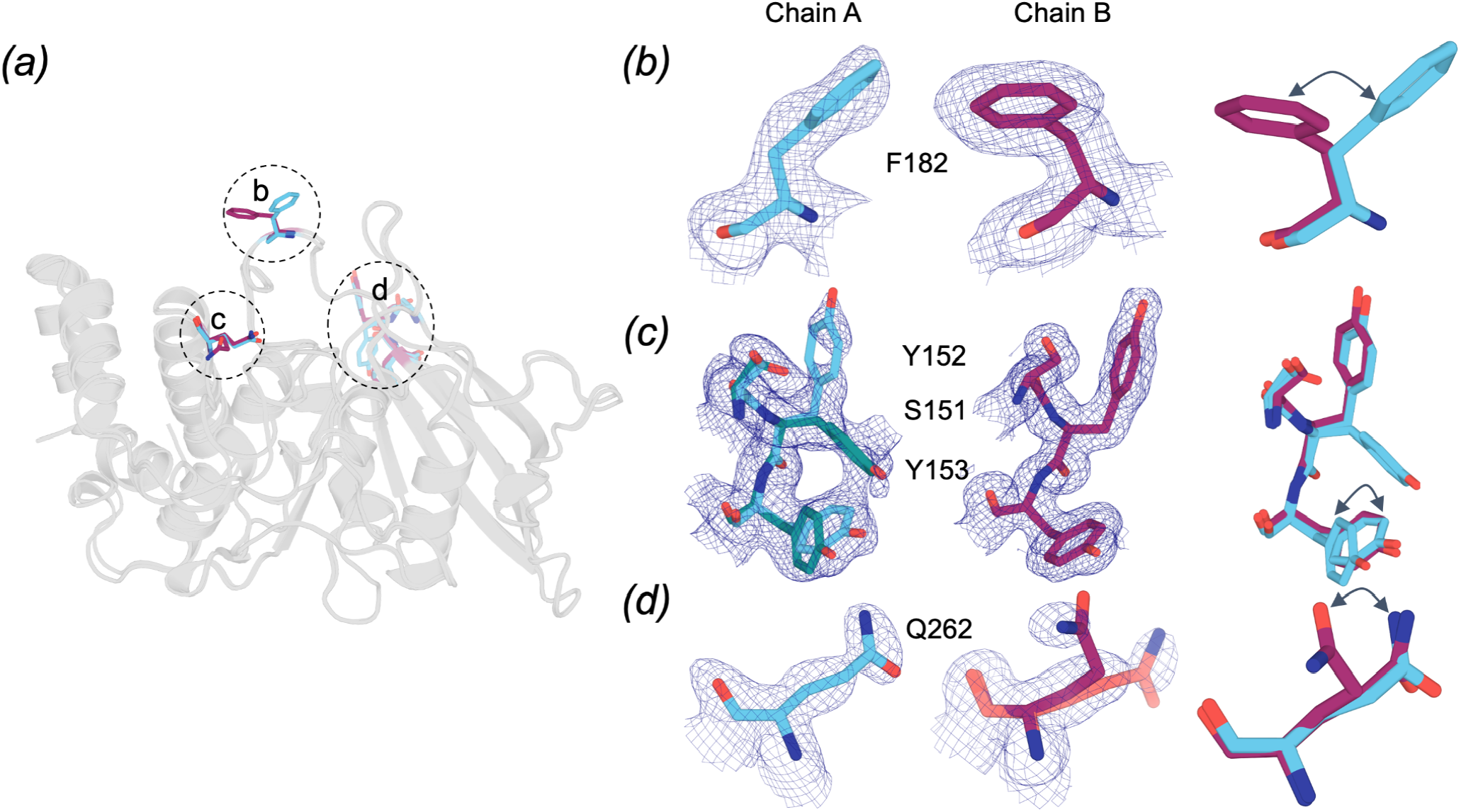
Examples of inter-chain differences in local alternate conformations. These examples demonstrate differences in side-chain conformations between chain A (cyan) and B (maroon).*(a)*Overlay of chains A and B, highlighting the locations of the residues displayed in panels *(b)*–*(d)*.*(b)*Residue F182 of the WPD loop adopts a different conformation in each chain (2Fo-Fc map in blue mesh, 1 σ).*(c)*The allosteric Loop 11 (Keedy et al. 2018) is flexible in chain A but more ordered in chain B (2Fo-Fc map in blue mesh, 0.5 σ; alternate conformation A in cyan, B in teal).*(d)*Q262 of the active-site Q loop adopts a single conformation in chain A but alternate conformations in chain B (2Fo-Fc map in blue mesh, 1 σ; alternate conformation A in maroon, B in salmon).

Second, a set of correlated alternate conformations can also be seen in Loop 11 (L11; residues 151–153). The conformations of residues in this loop have previously been shown to be correlated to the WPD loop and α7 helix (Keedy et al. 2018). In our structure, chain A adopts dual conformations for these residues (**Fig. 3 *(c)*, left panel**), whereas chain B adopts a single conformation (**Fig. 3 *(c)*, middle panel**).

Third, another residue from the active site — Q262 of the Q loop, which mediates hydrolysis of the phosphocysteine intermediate as part of the catalytic mechanism (Brandão, Hengge, and Johnson 2010) — samples two conformations only in chain B (**Fig. 3 *(d)*, middle panel**). This is in contrast to chain A where Q262 can only be modeled in a single conformation (**Fig. 3 *(d)*, left panel**).

For F182, the conformational difference can be attributed to a unique direct crystal contact in chain B. For Loop 11, the situation is similar, although the direct crystal contact only involves residues 151-152 from the loop, suggesting that the inter-chain differences for the other residues in the loop (**Fig. 3 *(c)***) may be due to conformational coupling in this allosteric region (Keedy et al. 2018). These local instances of crystal contacts in chain B but not in chain A are consistent with chain B exhibiting more crystal contacts and lower protein flexibility overall (**Table 2**). By contrast to these first two examples, Q262 is remote from any crystal contacts in either chain — suggesting that at least some of the conformational differences between the two chains arise from indirect or allosteric effects of the differential crystal packing.

### Diverse conformations of an allosteric region near the C-terminus

Of the ∼350 structures of PTP1B in the PDB, the majority exhibit a disordered and therefore unmodeled C-terminus. The ordering of the C-terminus is shown to be coupled, albeit only partially, to the conformation of the WPD loop and the allosteric L16 site (Keedy et al. 2018; Sharma, Ebrahim, and Keedy 2023; Skaist Mehlman et al. 2023). In a previously published closed-state structure of PTP1B, the C-terminus was modeled as ordered with the L16 site in the closed conformation (**Fig. 4 *(a)***) (Pedersen et al. 2004). This can be contrasted with an open-state structure of PTP1B bound to an allosteric inhibitor, in which the C-terminus is disordered and the L16 site is open (**Fig. 4 *(a)***) (Wiesmann et al. 2004).

**Figure 4:**
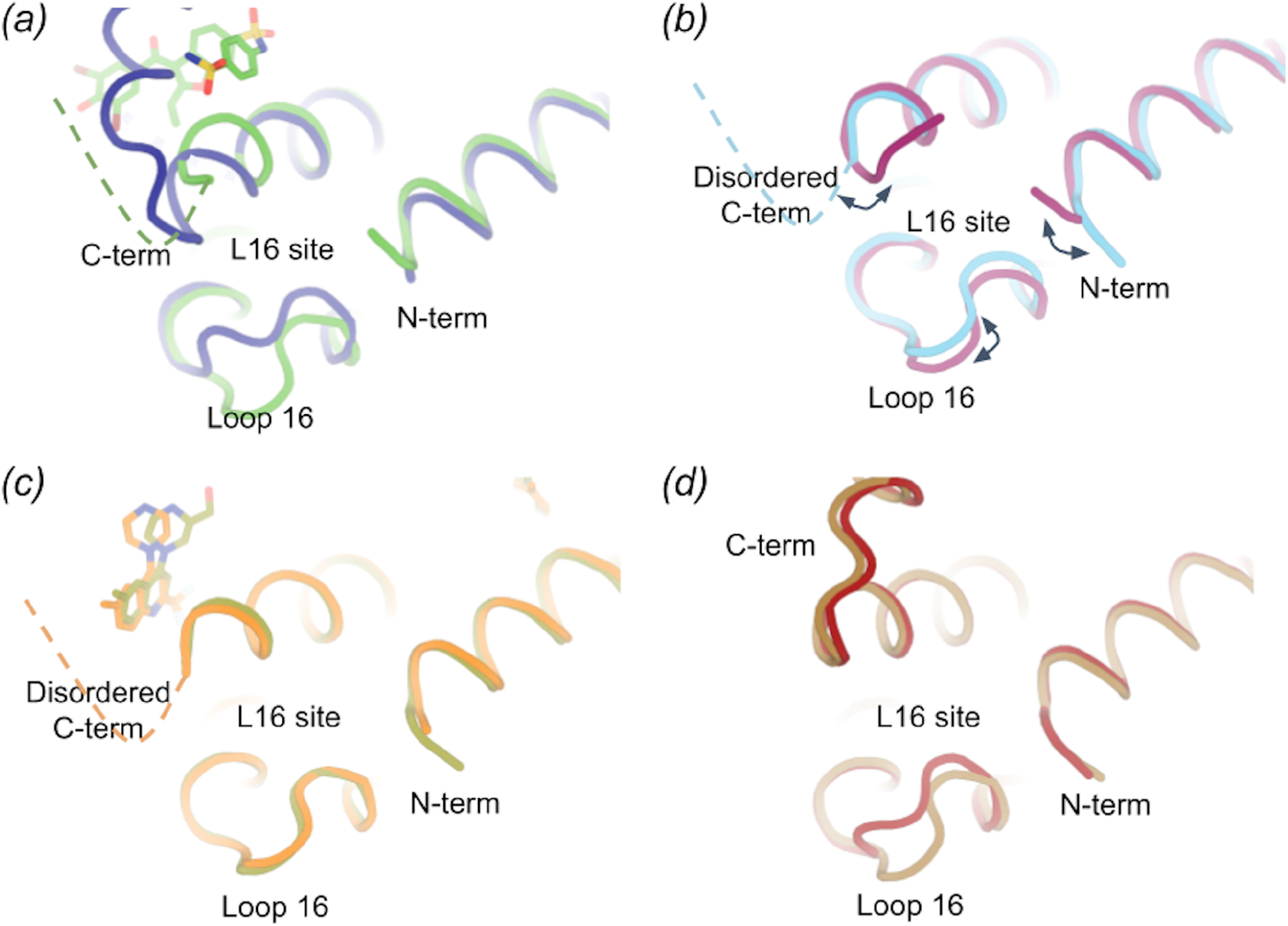
Unexpected differences in allosteric L16 site involving Loop 16, N-terminus, and C-terminus. *(a)*Comparison of PTP1B in the closed state (PDB ID: 1sug, dark blue) and in the open state bound to an allosteric inhibitor at the nearby BB site (PDB ID: 1t49, green) shows large changes in the regions constituting the allosteric L16 site.*(b)*An overlay of the two chains in our new structure (cyan: chain A, maroon: chain B) shows more subtle but significant differences in the three regions of the L16 site.*(c)*Two structures in the P 43 21 2 ligand-bound series (PDB IDs: 8g6a chain A, olive; 8g67 chain A, orange) (Greisman, Willmore, et al. 2023) show little to no changes in the three regions.*(d)*Two other structures in the P 43 21 2 ligand-bound series (PDB IDs: 8g69 chain A, dark red; 8g65 chain A, brown) (Greisman, Willmore, et al. 2023) show a uniquely reordered C-terminus as well as changes in Loop 16.

In our new high-resolution structure, a closer inspection of these regions shows disorder in the C-terminus in both chains, and a difference between chains in the L16 site involving a partial opening in chain B (∼1.4 Å Cα shift), coupled to a differently ordered N-terminus (**Fig. 4 *(b)***). The L16 site shift is notable because this loop typically exhibits bistable behavior, toggling between discrete open or closed states, but here exhibits an extra-open state. More generally, the changes seen between chains in our structure are notable in that they are smaller than, but reminiscent of, the changes observed between the canonical open and closed states of PTP1B (**Fig. 4 *(a)***).

A similar comparison with the recently reported ligand-bound P 43 21 2 structures indicates disorder in the C-terminus when two different small-molecule fragments bind in the nearby BB allosteric site (**Fig. 4 *(c)***), similar to what was observed with the BB3 allosteric inhibitor (**Fig. 4 *(a)***). In addition, in some other structures in this crystal form with other small-molecule fragments bound elsewhere in PTP1B, the C-terminus shows dramatic reordering into a non-helical conformation (**Fig. 4 *(d)***), in this case stabilized by extensive crystal contacts in only one of the two chains. Consistent with Cα distance analysis (**Fig. S1-S3**), in different of these structures L16 is modeled in either the open or the closed state (**Fig. 4 *(d)***). This is somewhat surprising given the previously reported correlation between an ordered C-terminal α7 helix and a closed L16 site (Keedy et al. 2018); it is possible that differently ordered C-terminus conformations have different allosteric effects on nearby regions. More generally, these observations are consistent with a previous report that the C-terminal α7 helix region can reorder into significantly distinct conformations, albeit involving adjacent bound ligands (Keedy et al. 2018), further suggesting that this key region of PTP1B is quite conformationally malleable.

### Unmodeled alternate conformations for activating mutations

Prior mutational analysis has provided information about the intramolecular interaction network in PTP1B, with certain sets of residues exhibiting evidence of coupling (Choy et al. 2017; Hjortness et al. 2018; Torgeson et al. 2022). One such analysis (Torgeson et al. 2022) demonstrated how double point mutations identified by coevolutionary sequence analysis (F225Y-R199N), distal from the active site, enhance catalysis by and reduce the stability of PTP1B. Upon closer inspection of the high-resolution F225Y-R199N crystal structure in the regions around the double mutations (**Fig. 5**), we observe evidence in the form of difference electron density patterns for unmodeled alternate conformations of several relevant residues, including both of the mutated residues themselves (F225Y, R199N) as well as several surrounding residues (including but not limited to F191, F174, L233, L204, C226) (**Fig. 5 *(b)***). Upon modeling the missing conformations, including distinct side-chain rotamers, subtler side-chain shifts within rotameric wells, and more complex movements of backbone plus side chains that are less straightforward to model, it becomes evident that these sites are more conformationally heterogeneous than originally modeled (**Fig. 5 *(c)***). Of particular note among these residues, F174 helps form the “floor” of the 197 allosteric site (Keedy et al. 2018), and F191 interacts directly with W179 of the active-site WPD loop. Our new observations here are consistent with the hypothesis based on NMR experiments that increased dynamics for the double mutant give rise to the increased activity and reduced stability (Torgeson et al. 2022), and provide atomistic insights into the conformational states that may be involved in those dynamics.

**Figure 5:**
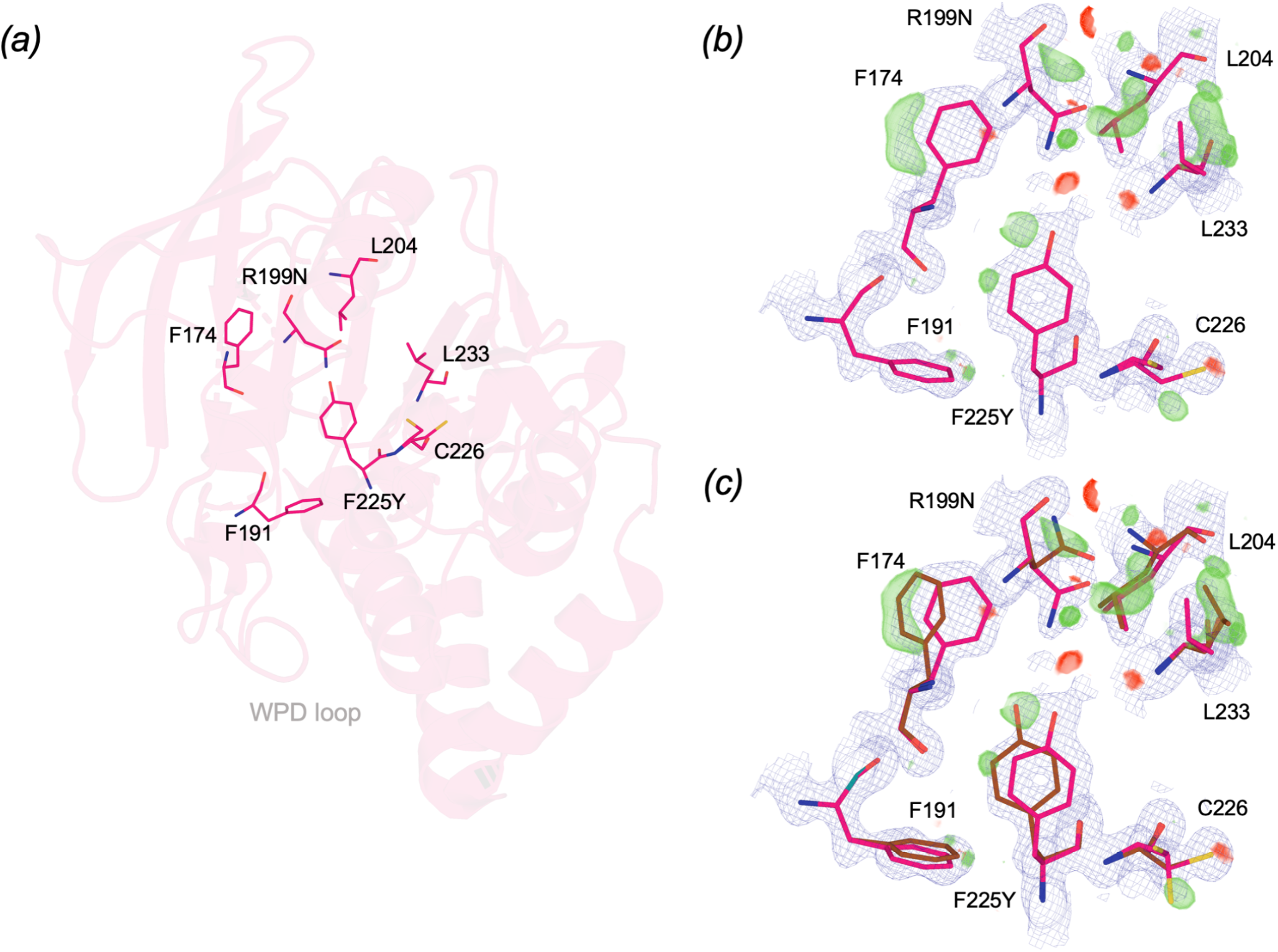
Modeling missing alternate conformations for a high-resolution double mutant structure. A cluster of residues centered around and including the point mutations F225Y and R199N in a high-resolution (1.24 Å) double mutant structure of PTP1B (PDB ID: 7mn9) (Torgeson et al. 2022) exhibits difference electron density features suggestive of unmodeled alternate conformations.Original conformations in pink; manually modeled alternate conformations in brown. *(a)*Zoomed-out view of residues displayed in subsequent panels. *(b)*Original model from 7mn9 (2Fo-Fc map in blue mesh, 1 σ; Fo-Fc map in green/red volume, +/-3 σ). *(c)*Updated model with manually modeled missing alternate conformations (2Fo-Fc map in blue mesh, 1 σ; Fo-Fc map in green/red meshes, +/-3 σ).

### Correlated conformations cluster near sites of mutations and ligands

Correlated motions between residues may convey allosteric information between distal sites in proteins such as PTP1B (Choy et al. 2017). We explored our multiconformer structure for any sites that may undergo such correlated motions, and identified two neighboring “sub-clusters” (**Fig. 6 *(a-b)***). Some residues in these sub-clusters (e.g. F191, T224) seem to exhibit compensating conformational heterogeneity, such that alternate conformations for one residue coincide with a single conformation for the other residue in one chain, with the situation reversed in the other chain.

**Figure 6:**
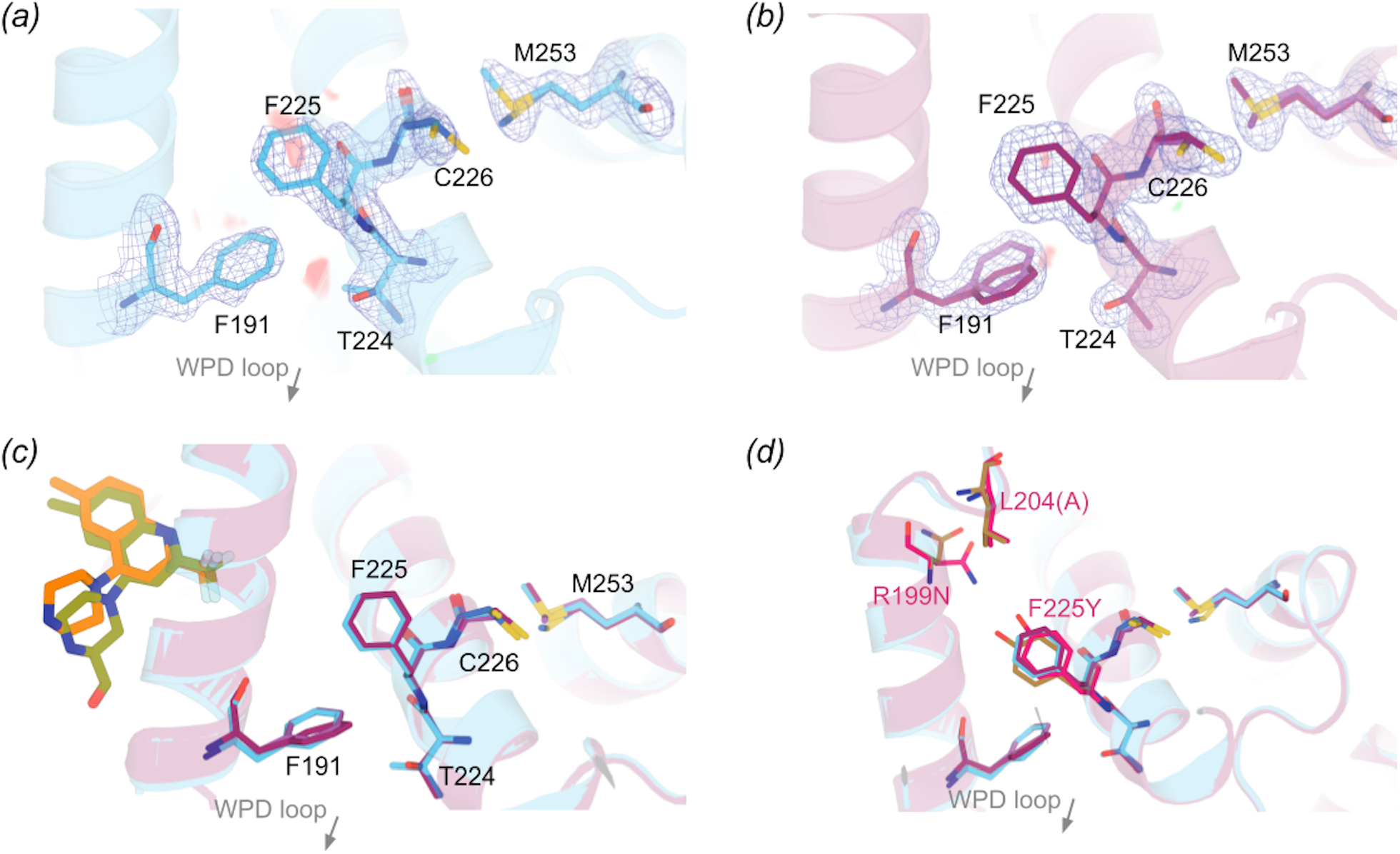
Clustered, correlated conformations localize to functionally linked and/or ligandable sites. Two sub-clusters of alternate conformations are identified, with one around the BB site (Wiesmann et al. 2004) and the other not previously reported in the allosteric network. Since these two sub-clusters are bridged by the functionally linked F225 (Torgeson et al. 2022), they cohere as a single coupled cluster.*(a-b)* Our new high-resolution apo WT structure (chain A, cyan; chain B, maroon) and electron density (2Fo-Fc map in blue mesh, 1 σ). We observe two adjacent sub-clusters with correlated alternate conformations. One involves the BB pathway (F191), and the other has not been previously reported as allosteric.*(c)*One of the two sub-clusters is also adjacent to recently reported small-molecule fragments that bind in the adjacent BB allosteric site, from the only structure series in the same crystal form as our structure (PDB IDs: 8g6a chain A, olive; 8g67 chain A, orange) (Greisman, Willmore, et al. 2023). *(d)*The two sub-clusters are also adjacent to several residues shown to influence activity upon mutation, including F225 which bridges the two sub-clusters (Torgeson et al. 2022). Here, we show an overlay of the F225Y-R199N double mutant structure (pink, PDB ID: 7mn9) with our WT structure. Additional remodeled conformations that were originally missing (see **Fig. 5**) are colored brown.

Interestingly, these sub-clusters lie adjacent to three regions of interest. The first is the catalytic WPD loop, including the eponymous W179 “anchor”, which directly contacts F191 — a residue which samples alternate conformations in one chain of our new structure (**Fig. 6 *(b)***). The second is the known BB allosteric site (Wiesmann et al. 2004; Greisman, Willmore, et al. 2023) (**Fig. 6 *(c)***). The third is a site of several recently reported activating point mutations (Torgeson et al. 2022) (**Fig. 6 *(d)***). These mutations include F225Y, which exhibits (previously unmodeled) conformational heterogeneity in the F225Y-R199N double mutant (**Fig. 5**). Thus, F225 serves as a “bridge” between the two sub-clusters we observe in the WT enzyme, which can thus be considered to form one cohesive cluster. This cluster collectively exhibits a combination of existing dynamics in WT PTP1B as well as the capacity for altered dynamics upon local perturbations that likely impact enzyme function.

## Discussion

In this study, we present the highest-resolution structure to date of apo WT PTP1B. This high resolution affords a detailed view of the conformational ensemble of this enzyme. Our structure is also the first of apo PTP1B in a recently discovered P 43 21 2 crystal form. For context, out of 350 total structures of PTP1B in the PDB, the representation of different crystal forms is far from uniform: there are 7 unique space groups, but 80% are in the P 31 2 1 space group. Based on our new apo structure as well as 5 other recent isomorphous ligand-bound structures (Greisman, Willmore, et al. 2023), the distinct packing in this crystal form appears to favor high-resolution diffraction, thereby helping to provide new windows into PTP1B’s conformational landscape.

Unlike most other PTP1B structures, our new structure’s unusual crystal form also accommodates two copies of the protein in distinct crystal packing environments. The distinct patterns of crystal contacts experienced by these two chains within the same asymmetric unit (**Table 2**) allowed us to perform controlled comparisons of the conformational ensemble of PTP1B, without the need to scale structure factors across datasets (Aldama, Dalton, and Hekstra 2023) or normalize B-factors across models (Ringe and Petsko 1986; Carugo and Argos 1997; Vihinen, Torkkila, and Riikonen 1994). Notably, the chain that was least impacted by crystal contacts exhibited noticeably higher disorder, with visually dispersed electron density, higher B-factors (**Table 2, Fig. 2**), and more alternate conformations in several regions (**Fig. 3**). Detailed inter-chain comparisons revealed that functional sites, such as the active site and allosteric sites, had the largest responses in terms of backbone shifts (**Fig. 1, Fig. 4**), side-chain rotamer changes (**Fig. 3**), and atomic disorder monitored by B-factors (**Fig. 2**). This localization of conformational flexibility to functional sites is consistent with the view that protein dynamics underlies and enables biological function (Tzeng and Kalodimos 2009; Fraser et al. 2009; Wei et al. 2016; Kim et al. 2017; Greisman, Dalton, et al. 2023).

In particular for PTP1B, we highlight a region centered on the α4 helix, including the flanking α3 and α5 helices, that exhibits apparently coupled conformational heterogeneity. Interestingly, this region lies adjacent to the previously identified BB allosteric site (Wiesmann et al. 2004) which also binds recently reported small-molecule fragments (Greisman, Willmore, et al. 2023) (**Fig. 6 *(d)***). Notably, several mutations in this area were recently reported to affect enzyme function, including F225Y and R199N which together increase catalysis and decrease stability (Torgeson et al. 2022). Our reanalysis of the previously reported high-resolution structure of the F225Y-R199N double mutant reveals compelling evidence in the electron density for missing, unmodeled alternate conformations for these two mutated residues as well as several neighboring residues (**Fig. 5**) — complementing our observations of conformational heterogeneity in this region of our new high-resolution WT structure. These crystallographic observations thus serve as unexpected additional validation of the previously proposed idea, based on NMR data and assuming a rigid crystal structure, that these mutations influence PTP1B function by imparting changes in protein dynamics (Torgeson et al. 2022). These results are also consistent with the notion that WT PTP1B may harbor latent dynamic wiring that can be modulated by mutations to alter function (Tokuriki and Tawfik 2009).

The alternate conformations for the double mutant mentioned above were especially identifiable due to the structure’s particularly high resolution: indeed, it has the highest resolution of all 350 available PTP1B structures (1.24 Å). However, unmodeled alternate conformations are surprisingly common in crystal structures across the PDB (Lang et al. 2010; Riley et al. 2021; Wankowicz et al. 2023; Stachowski and Fischer 2023). If properly modeled, they could likely help explain functional effects of mutations and ligands (Wankowicz et al. 2022), allosteric mechanisms, and other phenomena for PTP1B and other systems. Moreover, other biophysical perturbations (Keedy 2019) such as variable temperature (Fraser et al. 2011; Keedy et al. 2014, 2018; Fischer 2021; Stachowski et al. 2022; Sharma, Ebrahim, and Keedy 2023; Skaist Mehlman et al. 2023; Greisman, Dalton, et al. 2023; Thorne 2023) or pressure (Urayama, Phillips, and Gruner 2002; Guerrero et al. 2023) could reveal additional, previously “hidden” conformational heterogeneity or excited states that could provide further insights into biological mechanisms for PTPs as well as other proteins. These areas represent promising avenues for future study.

## Materials and Methods

### Protein expression and purification

For these experiments, PTP1B (residues 1-321) with an additional C-terminal His tag was expressed. Briefly, *E. coli* BL21 cells were transformed with 10 ng of His-tagged pET21b - human PTP1B (1-321) plasmid and plated on the Luria broth (LB) medium agar plate with ampicillin and incubated overnight at 37°C. A single bacterial colony was picked and grown overnight in LB media with ampicillin at 37°C in a shaker incubator as the primary culture. The next day, the required amount of primary culture was added to fresh LB media and allowed to grow at 37°C. Once the optical density (OD) value reaches 0.4–0.6, 1 mM IPTG was added and the culture was grown overnight in a shaker incubator at 200°C. The culture was pelleted by centrifugation at 4000 rpm for 1 hr. Pellets were lysed immediately or stored at -20°C.

Bacterially expressed His-PTP1B (residues 1-321) was purified by Ni-NTA (nitrilotriacetic acid). Briefly, the bacterial pellet was solubilized in lysis buffer (20 mM NaH_2_PO_4_, 300 mM NaCl, 1 mM TCEP, protease inhibitor cocktail tablet, pH 8) and lysed using a sonicator on ice (amplitude 40%, pulse on: 5 seconds, pulse off: 10 seconds, total 10 min). During sonication, a Ni-NTA column was equilibrated with lysis buffer. Lysate was centrifuged at 4000 rpm, 40°C for 1 hr and supernatant was added to the pre equilibrated Ni-NTA column and incubated for 1 hr on a clinical rotor at 40°C. After incubation, the column was washed with 5 volumes of wash buffer (20 mM NaH_2_PO_4_, 300 mM NaCl, 1 mM TCEP, 20 mM imidazole, pH 8) and finally eluted with elution buffer (20 mM NaH_2_PO_4_, 300 mM NaCl, 1 mM TCEP, 250mM imidazole, pH 8). Imidazole from the protein was removed using a Zeba spin buffer exchange column (ThermoFisher Scientific, model # 89882) and PTP1B was stored in a 5 mM TCEP solution (50 mM HEPES, 150 mM NaCl, 5 mM TCEP, pH 8) at 40°C. Protein quantification was performed using the Bradford method. Purified PTP1B was then buffer exchanged (50 mM HEPES, 150 mM NaCl, pH 8) to remove TCEP before crystallization.

### Crystallization

WT PTP1B at stock concentration of 1 mM (final concentration of 0.30 mM) was incubated with water-soluble cholesterol at stock concentration of 0.1 M (final concentration of 9.1 mM) in storage buffer (10 mM Tris pH 7.5, 0.2 mM EDTA, 25 mM NaCl, 3 mM DTT) for 3 hours at room temperature. This mixture was used to set up crystallization sitting drops in 96-well low-profile Art Robbins INTELLI-PLATE trays using an SPT Labtech Mosquito Xtal3, in a ratio of 0.1 μL protein + 0.1μL well solution (0.1M MgCl_2_, 0.1 M Hepes pH 7.0, 15% w/v PEG 4000; from ProComplex commercial screen), which were then incubated at 4°C. Crystals grew within 5–7 days to a final size of 25–50 μm.

### X-ray data collection

Crystals were harvested using MiTeGen microloops of appropriate size, then cryocooled by hand-plunging into liquid nitrogen. Diffraction data were collected remotely at the NYX beamline at the National Synchrotron Light Source II (NSLS-II). Single crystals were exposed to X-rays under a continuous cryostream (100 K), with 0.2° crystal rotation and 0.12 seconds exposure per image, for a total of 360° across 1800 images.

### Crystallographic data processing and modeling

Diffraction data was processed using the DIALS pipeline (Winter et al. 2022) via xia2 (G. Winter 2009). All 1800 images (360°) were included in processing, and the space group P 43 21 2 was enforced. The resolution limit was automatically selected by DIALS (Winter et al. 2022). The resulting data processing statistics were favorable (**Table 1**).

The resulting merged structure factors file was used for molecular replacement with Phaser (McCoy et al. 2007) with PDB ID 1t49 as the search model. We also ran the program Xtriage (Zwart,Grosse-Kunstleve, and Adams 2005) to obtain the Matthew’s coefficient (**Table 1**). The Matthew’s coefficient of 2.21 indicated the need to place two copies of the protein molecule in the asymmetric unit (non-identical copies). The placement of the two chains in the asymmetric unit was chosen to match that of the other recently published structures in this crystal form (Greisman, Willmore, et al. 2023), for consistency and ease of interpretation.

Iterative modeling with Coot (Emsley et al. 2010) between rounds of refinement was performed. Several iterations of refinement were performed using the phenix.refine program within the PHENIX suite (Liebschner et al. 2019). Refinement was performed with the ‘anisotropic B factor’ flag and ‘update waters’ set to true. ‘Update waters’ was turned off in later refinement rounds as refinement approached convergence. By default, non-crystallographic symmetry between chains was not imposed. The resulting refinement statistics were favorable (**Table 1**).

### Analysis of models

The number of crystal contacts and total crystal surface area in **Table 2** was calculated using the PISA web server (Krissinel and Henrick 2007).

The average B-factor values for each chain in **Table 2** were calculated using Excel.

Per-residue Cα distances between chains were calculated using VMD (Humphrey, Dalke, and Schulten 1996) using the following steps:

- File → New Molecule
- Extensions → Analysis → MultiSeq
- Tools → Stamp Structural Alignment
- Export results

The spectrum bars representing the ranges of Cα distance and B-factor values were created using the spectrumbar.py script from PyMol Wiki. The colors used for the spectrum bar are (in order) blue, cyan, green, and yellow, with rectangular ends, a radius of 1.5, and a length of 50.0.

Coloring the structures by Cα distance was done using the colorbyrmsd.py script from PyMol Wiki. All structure figures were generated using PyMol (Schrödinger and DeLano 2020).

## Data availability

Model, structure factor, and map files are available in the Protein Data Bank using PDB ID (accession code) 8u1e. Raw X-ray diffraction images for this dataset have also been deposited at proteindiffraction.org.

## Acknowledgments

DAK is supported by NIH R35 GM133769.

We thank Kevin Battaile and the staff at NYX for assistance with data collection.

## References

Ala, Paul J., Lucie Gonneville, Milton C. Hillman, Mary Becker-Pasha, Min Wei, Brian G. Reid, Ronald Klabe, et al. 2006. “Structural Basis for Inhibition of Protein-Tyrosine Phosphatase 1B by Isothiazolidinone Heterocyclic Phosphonate Mimetics.” The Journal of Biological Chemistry 281 (43): 32784–95.

Aldama, Luis A., Kevin M. Dalton, and Doeke R. Hekstra. 2023. “Correcting Systematic Errors in Diffraction Data with Modern Scaling Algorithms.” Acta Crystallographica. Section D, Structural Biology, September. 10.1107/S2059798323005776.

Barstow, Buz, Nozomi Ando, Chae Un Kim, and Sol M. Gruner. 2008. “Alteration of Citrine Structure by Hydrostatic Pressure Explains the Accompanying Spectral Shift.” Proceedings of the National Academy of Sciences of the United States of America 105 (36): 13362–66.

Bedem, Henry van den, Gira Bhabha, Kun Yang, Peter E. Wright, and James S. Fraser. 2013. “Automated Identification of Functional Dynamic Contact Networks from X-Ray Crystallography.” Nature Methods 10 (9): 896–902.

Berman, Helen M., John Westbrook, Zukang Feng, Gary Gilliland, T. N. Bhat, Helge Weissig, Ilya N. Shindyalov, and Philip E. Bourne. 2000. “The Protein Data Bank.” Nucleic Acids Research 28 (1): 235–42.

Bhabha, Gira, Justin T. Biel, and James S. Fraser. 2015. “Keep on Moving: Discovering and Perturbing the Conformational Dynamics of Enzymes.” Accounts of Chemical Research 48 (2): 423–30.

Bhattacharyya, Moitrayee, Soma Ghosh, and Saraswathi Vishveshwara. 2016. “Protein Structure and Function: Looking through the Network of Side-Chain Interactions.” Current Protein & Peptide Science 17 (1): 4–25.

Brandão, Tiago A. S., Alvan C. Hengge, and Sean J. Johnson. 2010. “Insights into the Reaction of Protein-Tyrosine Phosphatase 1B: Crystal Structures for Transition State Analogs of Both Catalytic Steps.” The Journal of Biological Chemistry 285 (21): 15874–83.

Carugo, O., and P. Argos. 1997. “Protein-Protein Crystal-Packing Contacts.” Protein Science: A Publication of the Protein Society 6 (10): 2261–63.

Choy, Meng S., Yang Li, Luciana E. S. F. Machado, Micha B. A. Kunze, Christopher R. Connors, Xingyu Wei, Kresten Lindorff-Larsen, Rebecca Page, and Wolfgang Peti. 2017. “Conformational Rigidity and Protein Dynamics at Distinct Timescales Regulate PTP1B Activity and Allostery.” Molecular Cell 65 (4): 644–58.e5.

Davis, Ian W., W. Bryan Arendall 3rd, David C. Richardson, and Jane S. Richardson. 2006. “The Backrub Motion: How Protein Backbone Shrugs When a Sidechain Dances.” Structure 14 (2): 265–74.

Deis, Lindsay N., Charles W. Pemble 4th, Yang Qi, Andrew Hagarman, David C. Richardson, Jane S. Richardson, and Terrence G. Oas. 2014. “Multiscale Conformational Heterogeneity in Staphylococcal Protein a: Possible Determinant of Functional Plasticity.” Structure 22 (10): 1467–77.

Emsley, P., B. Lohkamp, W. G. Scott, and K. Cowtan. 2010. “Features and Development of Coot.” Acta Crystallographica. Section D, Biological Crystallography 66 (Pt 4): 486–501.

Fischer, Marcus. 2021. “Macromolecular Room Temperature Crystallography.” Quarterly Reviews of Biophysics 54 (January): e1.

Fischer, Marcus, Brian K. Shoichet, and James S. Fraser. 2015. “One Crystal, Two Temperatures: Cryocooling Penalties Alter Ligand Binding to Transient Protein Sites.” Chembiochem: A European Journal of Chemical Biology 16 (11): 1560–64.

Fraser, James S., Henry van den Bedem, Avi J. Samelson, P. Therese Lang, James M. Holton, Nathaniel Echols, and Tom Alber. 2011. “Accessing Protein Conformational Ensembles Using Room-Temperature X-Ray Crystallography.” Proceedings of the National Academy of Sciences of the United States of America 108 (39): 16247–52.

Fraser, James S., Michael W. Clarkson, Sheena C. Degnan, Renske Erion, Dorothee Kern, and Tom Alber. 2009. “Hidden Alternative Structures of Proline Isomerase Essential for Catalysis.” Nature 462 (7273): 669–73.

Greisman, Jack B., Kevin M. Dalton, Dennis E. Brookner, Margaret A. Klureza, Candice J. Sheehan, In-Sik Kim, Robert W. Henning, Silvia Russi, and Doeke R. Hekstra. 2023. “Resolving Conformational Changes That Mediate a Two-Step Catalytic Mechanism in a Model Enzyme.” bioRxiv : The Preprint Server for Biology, June. 10.1101/2023.06.02.543507.

Greisman, Jack B., Lindsay Willmore, Christine Y. Yeh, Fabrizio Giordanetto, Sahar Shahamadtar, Hunter Nisonoff, Paul Maragakis, and David E. Shaw. 2023. “Discovery and Validation of the Binding Poses of Allosteric Fragment Hits to Protein Tyrosine Phosphatase 1b: From Molecular Dynamics Simulations to X-Ray Crystallography.” Journal of Chemical Information and Modeling 63 (9): 2644–50.

Guerrero, Liliana, Ali Ebrahim, Blake T. Riley, Minyoung Kim, Qingqiu Huang, Aaron D. Finke, and Daniel A. Keedy. 2023. “Pushed to Extremes: Distinct Effects of High Temperature vs. Pressure on the Structure of an Atypical Phosphatase.” bioRxiv : The Preprint Server for Biology, May. 10.1101/2023.05.02.538097.

Hjortness, Michael K., Laura Riccardi, Akarawin Hongdusit, Peter H. Zwart, Banumathi Sankaran, Marco De Vivo, and Jerome M. Fox. 2018. “Evolutionarily Conserved Allosteric Communication in Protein Tyrosine Phosphatases.” Biochemistry 57 (45): 6443–51.

Humphrey, W., A. Dalke, and K. Schulten. 1996. “VMD-Visual Molecular Dynamics J Mol Graph 14: 33--38.”

Jacobson, Matthew P., Richard A. Friesner, Zhexin Xiang, and Barry Honig. 2002. “On the Role of the Crystal Environment in Determining Protein Side-Chain Conformations.” Journal of Molecular Biology 320 (3): 597–608.

Johansson, Kristoffer E., and Kresten Lindorff-Larsen. 2018. “Structural Heterogeneity and Dynamics in Protein Evolution and Design.” Current Opinion in Structural Biology 48 (February): 157–63.

Keedy, Daniel A. 2019. “Journey to the Center of the Protein: Allostery from Multitemperature Multiconformer X-Ray Crystallography.” Acta Crystallographica. Section D, Structural Biology 75 (Pt 2): 123–37.

Keedy, Daniel A., Henry van den Bedem, David A. Sivak, Gregory A. Petsko, Dagmar Ringe, Mark A. Wilson, and James S. Fraser. 2014. “Crystal Cryocooling Distorts Conformational Heterogeneity in a Model Michaelis Complex of DHFR.” Structure 22 (6): 899–910.

Keedy, Daniel A., James S. Fraser, and Henry van den Bedem. 2015. “Exposing Hidden Alternative Backbone Conformations in X-Ray Crystallography Using qFit.” PLoS Computational Biology 11 (10): e1004507.

Keedy, Daniel A., Zachary B. Hill, Justin T. Biel, Emily Kang, T. Justin Rettenmaier, José Brandão-Neto, Nicholas M. Pearce, Frank von Delft, James A. Wells, and James S. Fraser. 2018. “An Expanded Allosteric Network in PTP1B by Multitemperature Crystallography, Fragment Screening, and Covalent Tethering.” eLife 7 (June). 10.7554/eLife.36307.

Kim, Tae Hun, Pedram Mehrabi, Zhong Ren, Adnan Sljoka, Christopher Ing, Alexandr Bezginov, Libin Ye, Régis Pomès, R. Scott Prosser, and Emil F. Pai. 2017. “The Role of Dimer Asymmetry and Protomer Dynamics in Enzyme Catalysis.” Science 355 (6322). 10.1126/science.aag2355.

Krissinel, Evgeny, and Kim Henrick. 2007. “Inference of Macromolecular Assemblies from Crystalline State.” Journal of Molecular Biology 372 (3): 774–97.

Lang, P. Therese, Ho-Leung Ng, James S. Fraser, Jacob E. Corn, Nathaniel Echols, Mark Sales, James M. Holton, and Tom Alber. 2010. “Automated Electron-Density Sampling Reveals Widespread Conformational Polymorphism in Proteins.” Protein Science: A Publication of the Protein Society 19 (7): 1420–31.

Liebschner, Dorothee, Pavel V. Afonine, Matthew L. Baker, Gábor Bunkóczi, Vincent B. Chen, Tristan Croll, Bradley Hintze, et al. 2019. “Macromolecular Structure Determination Using X-Rays, Neutrons and Electrons: Recent Developments in Phenix.” Acta Crystallographica. Section D, Structural Biology 75 (Pt 10): 861–77.

Lovell, S. C., J. M. Word, J. S. Richardson, and D. C. Richardson. 2000. “The Penultimate Rotamer Library.” Proteins 40 (3): 389–408.

Martin, Alberto J. M., Michele Vidotto, Filippo Boscariol, Tomàs Di Domenico, Ian Walsh, and Silvio C. E. Tosatto. 2011. “RING: Networking Interacting Residues, Evolutionary Information and Energetics in Protein Structures.” Bioinformatics 27 (14): 2003–5.

McCoy, Airlie J., Ralf W. Grosse-Kunstleve, Paul D. Adams, Martyn D. Winn, Laurent C. Storoni, and Randy J. Read. 2007. “Phaser Crystallographic Software.” Journal of Applied Crystallography 40 (Pt 4): 658–74.

Morris, Rhiannon, Narelle Keating, Cyrus Tan, Hao Chen, Artem Laktyushin, Tamanna Saiyed, Nicholas P. D. Liau, et al. 2023. “Structure Guided Studies of the Interaction between PTP1B and JAK.” Communications Biology 6 (1): 641.

Pedersen, Anja K., G. Ünther H. Peters G, Karin B. Møller, Lars F. Iversen, and Jette S. Kastrup. 2004. “Water-Molecule Network and Active-Site Flexibility of Apo Protein Tyrosine Phosphatase 1B.” Acta Crystallographica. Section D, Biological Crystallography 60 (Pt 9): 1527–34.

Riley, Blake T., Stephanie A. Wankowicz, Saulo H. P. de Oliveira, Gydo C. P. van Zundert, Daniel W. Hogan, James S. Fraser, Daniel A. Keedy, and Henry van den Bedem. 2021. “qFit 3: Protein and Ligand Multiconformer Modeling for X-Ray Crystallographic and Single-Particle Cryo-EM Density Maps.” Protein Science: A Publication of the Protein Society 30 (1): 270–85.

Ringe, D., and G. A. Petsko. 1986. “Study of Protein Dynamics by X-Ray Diffraction.” Methods in Enzymology 131 (January): 389–433.

Schrödinger, L., and Warren DeLano. 2020. “PyMOL.”

Sharma, Shivani, Ali Ebrahim, and Daniel A. Keedy. 2023. “Room-Temperature Serial Synchrotron Crystallography of the Human Phosphatase PTP1B.” Acta Crystallographica. Section F, Structural Biology and Crystallization Communications 79 (Pt 1): 23–30.

Skaist Mehlman, Tamar, Justin T. Biel, Syeda Maryam Azeem, Elliot R. Nelson, Sakib Hossain, Louise Dunnett, Neil G. Paterson, et al. 2023. “Room-Temperature Crystallography Reveals Altered Binding of Small-Molecule Fragments to PTP1B.” eLife 12 (March). 10.7554/eLife.84632.

Stachowski, Timothy R., and Marcus Fischer. 2023. “FLEXR: Automated Multi-Conformer Model Building Using Electron-Density Map Sampling.” Acta Crystallographica. Section D, Structural Biology 79 (Pt 5): 354–67.

Stachowski, Timothy R., Murugendra Vanarotti, Jayaraman Seetharaman, Karlo Lopez, and Marcus Fischer. 2022. “Water Networks Repopulate Protein-Ligand Interfaces with Temperature.” Angewandte Chemie 61 (31): e202112919.

Thorne, Robert E. 2023. “Determining Biomolecular Structures near Room Temperature Using X-Ray Crystallography: Concepts, Methods and Future Optimization.” Acta Crystallographica. Section D, Structural Biology 79 (Pt 1): 78–94.

Tokuriki, Nobuhiko, and Dan S. Tawfik. 2009. “Stability Effects of Mutations and Protein Evolvability.” Current Opinion in Structural Biology 19 (5): 596–604.

Torgeson, Kristiane R., Michael W. Clarkson, Daniele Granata, Kresten Lindorff-Larsen, Rebecca Page, and Wolfgang Peti. 2022. “Conserved Conformational Dynamics Determine Enzyme Activity.” Science Advances 8 (31): eabo5546.

Tyka, Michael D., Daniel A. Keedy, Ingemar André, Frank Dimaio, Yifan Song, David C. Richardson, Jane S. Richardson, and David Baker. 2011. “Alternate States of Proteins Revealed by Detailed Energy Landscape Mapping.” Journal of Molecular Biology 405 (2): 607–18.

Tzeng, Shiou-Ru, and Charalampos G. Kalodimos. 2009. “Dynamic Activation of an Allosteric Regulatory Protein.” Nature 462 (7271): 368–72.

Urayama, Paul, George N. Phillips Jr, and Sol M. Gruner. 2002. “Probing Substates in Sperm Whale Myoglobin Using High-Pressure Crystallography.” Structure 10 (1): 51–60.

Vihinen, M., E. Torkkila, and P. Riikonen. 1994. “Accuracy of Protein Flexibility Predictions.” Proteins 19 (2): 141–49.

Wankowicz, Stephanie A., Saulo H. de Oliveira, Daniel W. Hogan, Henry van den Bedem, and James S. Fraser. 2022. “Ligand Binding Remodels Protein Side-Chain Conformational Heterogeneity.” eLife 11 (March). 10.7554/eLife.74114.

Wankowicz, Stephanie A., Ashraya Ravikumar, Shivani Sharma, Blake T. Riley, Akshay Raju, Daniel W. Hogan, Henry van den Bedem, Daniel A. Keedy, and James S. Fraser. 2023. “Uncovering Protein Ensembles: Automated Multiconformer Model Building for X-Ray Crystallography and Cryo-EM.” bioRxiv : The Preprint Server for Biology, June. 10.1101/2023.06.28.546963.

Wei, Guanghong, Wenhui Xi, Ruth Nussinov, and Buyong Ma. 2016. “Protein Ensembles: How Does Nature Harness Thermodynamic Fluctuations for Life? The Diverse Functional Roles of Conformational Ensembles in the Cell.” Chemical Reviews 116 (11): 6516–51.

Whittier, Sean K., Alvan C. Hengge, and J. Patrick Loria. 2013. “Conformational Motions Regulate Phosphoryl Transfer in Related Protein Tyrosine Phosphatases.” Science 341 (6148): 899–903.

Wiesmann, Christian, Kenneth J. Barr, Jenny Kung, Jiang Zhu, Daniel A. Erlanson, Wang Shen, Bruce J. Fahr, et al. 2004. “Allosteric Inhibition of Protein Tyrosine Phosphatase 1B.” Nature Structural & Molecular Biology 11 (8): 730–37.

Winter, G. 2009. “xia2: An Expert System for Macromolecular Crystallography Data Reduction.” Journal of Applied Crystallography 43 (1): 186–90.

Winter, Graeme, James Beilsten-Edmands, Nicholas Devenish, Markus Gerstel, Richard J. Gildea, David McDonagh, Elena Pascal, David G. Waterman, Benjamin H. Williams, and Gwyndaf Evans. 2022. “DIALS as a Toolkit.” Protein Science: A Publication of the Protein Society 31 (1): 232–50.

Yabukarski, Filip, Justin T. Biel, Margaux M. Pinney, Tzanko Doukov, Alexander S. Powers, James S. Fraser, and Daniel Herschlag. 2020. “Assessment of Enzyme Active Site Positioning and Tests of Catalytic Mechanisms through X-Ray-Derived Conformational Ensembles.” Proceedings of the National Academy of Sciences of the United States of America 117 (52): 33204–15.

Yeh, Christine Y., Jesus A. Izaguirre, Jack B. Greisman, Lindsay Willmore, Paul Maragakis, and David E. Shaw. 2023. “A Conserved Local Structural Motif Controls the Kinetics of PTP1B Catalysis.” bioRxiv. 10.1101/2023.02.28.529746.

Zwart, P. H., R. W. Grosse-Kunstleve, and P. D. Adams. 2005. “Xtriage and Fest: Automatic Assessment of X-Ray Data and Substructure Structure Factor Estimation.” CCP4 Newsletter, 2005.

