## Supplemental Figures for "High-resolution double vision of the archetypal protein tyrosine phosphatase"

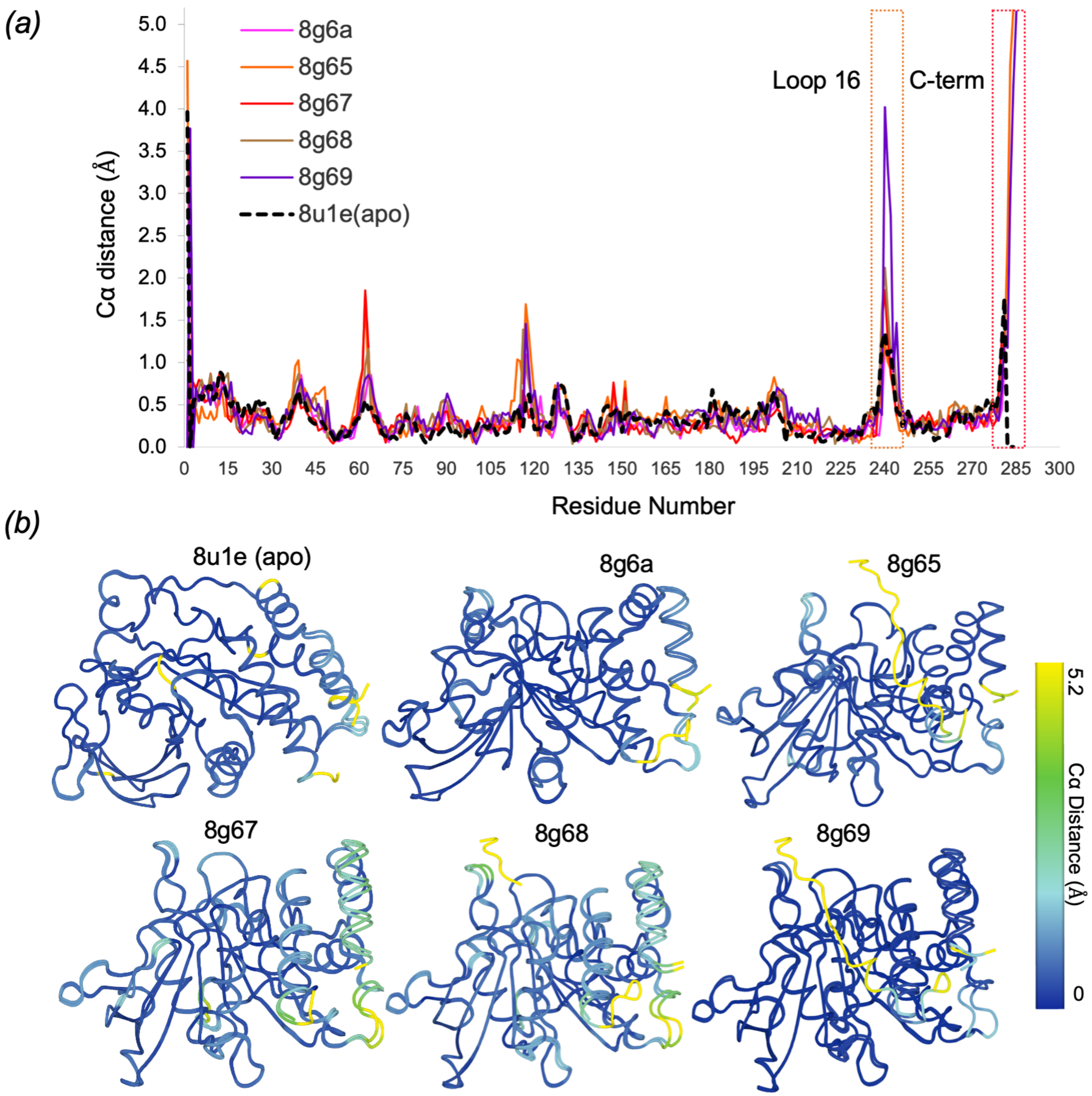


**Figure S1: Backbone displacements between chains across the P 43 21 2 series.**
Regions with highest backbone variability between the two chains are highlighted with colored dotted outlines for our high-resolution apo structure and the isomorphous ligand-bound structure series (Greisman, Willmore, et al. 2023).
*(a)* Plot of inter-chain Cα distance vs. amino acid sequence.
*(b)* Overlay of both chains with residues colored by inter-chain Cα distance.

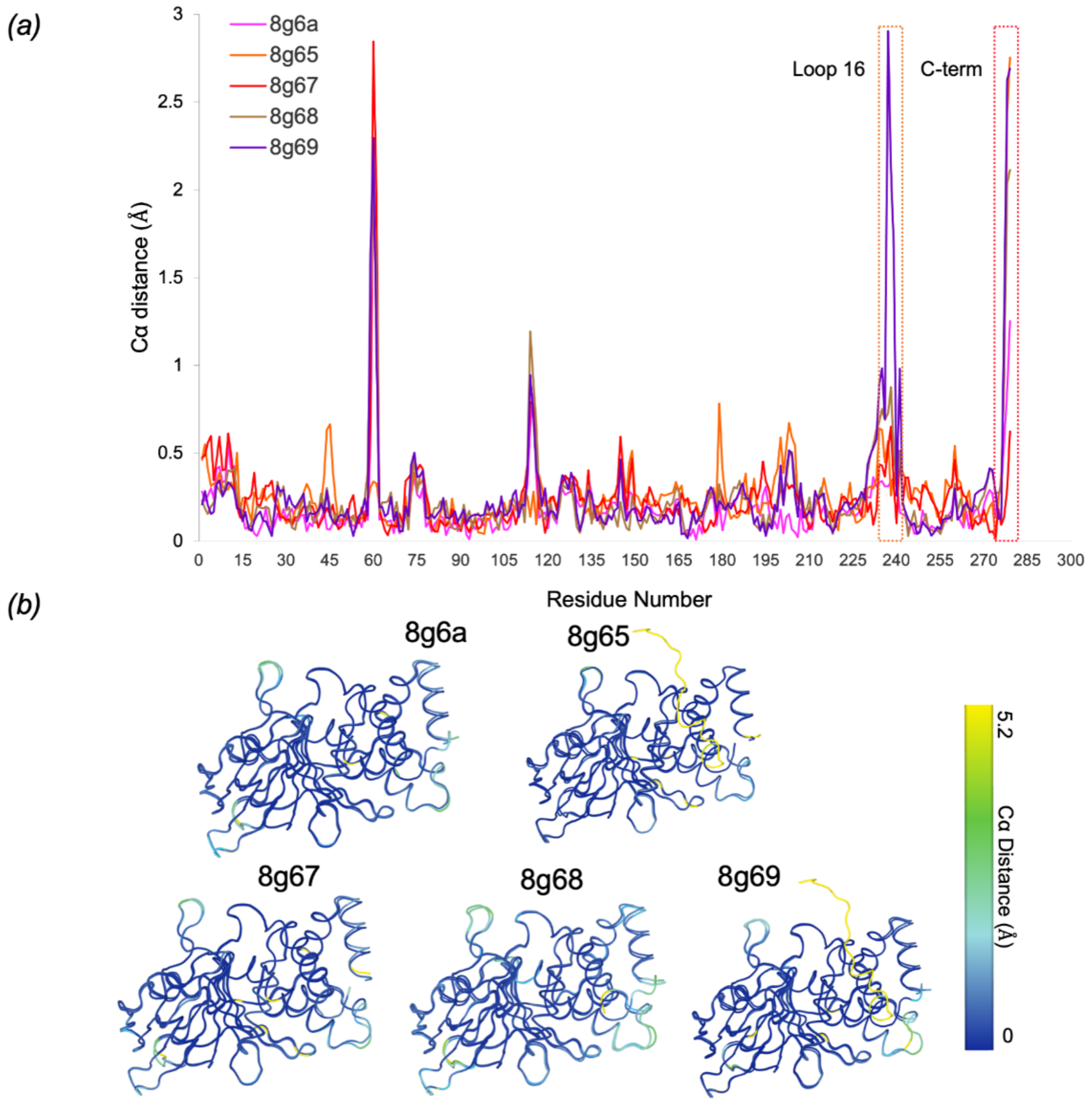


**Figure S2: Backbone displacements between chain A of 8u1e (apo) and of the ligand-bound P 43 21 2 series.**

Regions with highest backbone variability between structures for chain A are highlighted with colored dotted outlines for our high-resolution apo structure and the isomorphous ligand-bound structure series (Greisman, Willmore, et al. 2023).

*(a)* Plot of inter-structure (8u1e as reference) Cα distance vs. amino acid sequence.

*(b)* Overlay of chain A in 8u1e and the compared structure with residues colored by Cα distance.

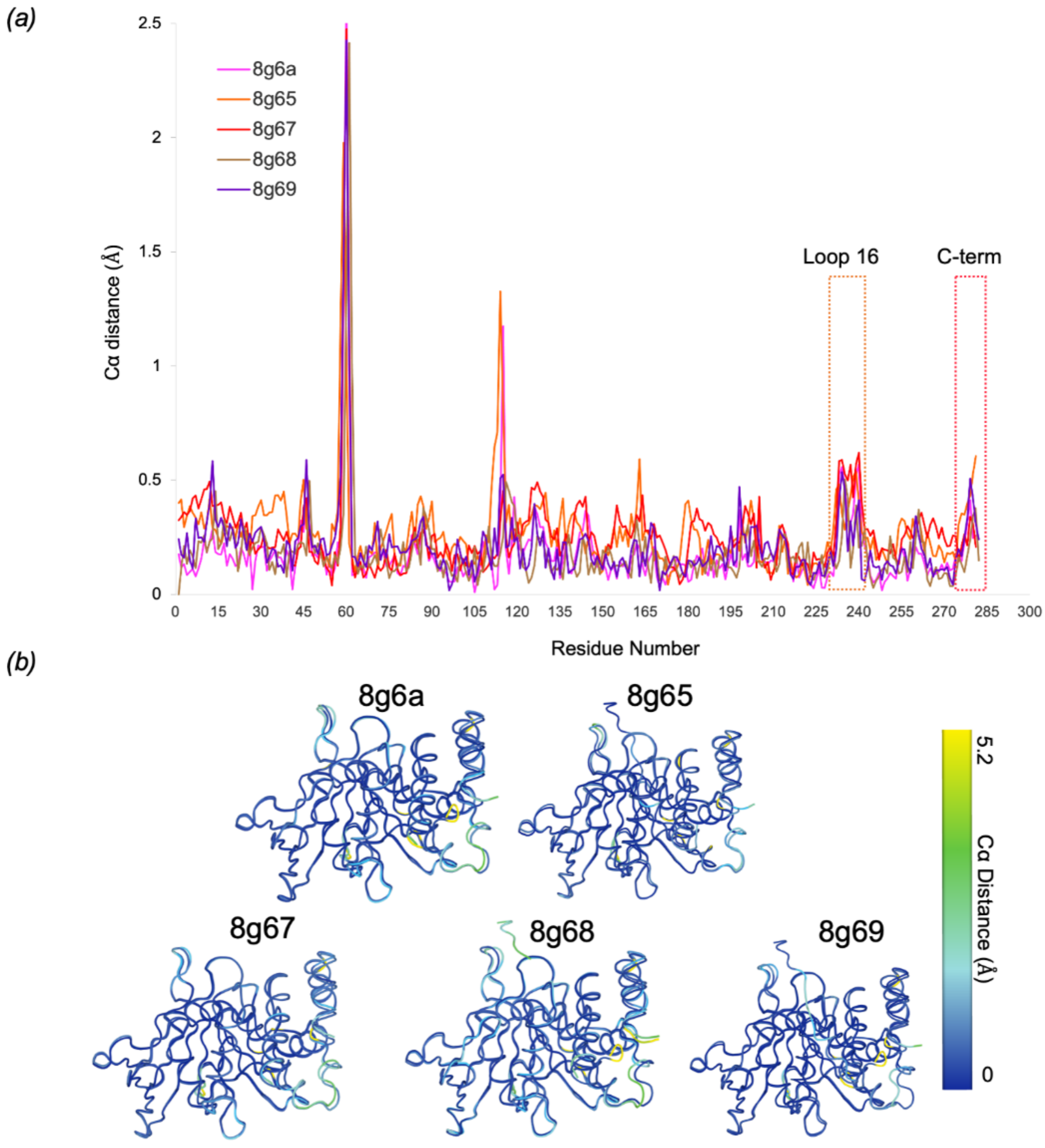


**Figure S3: Backbone displacements between chain B of 8u1e (apo) and of the ligand-bound P 43 21 2 series.**

Regions with highest backbone variability between structures for chain B are highlighted with colored dotted outlines for our high-resolution apo structure and the isomorphous ligand-bound structure series (Greisman, Willmore, et al. 2023).

*(a)* Plot of inter-structure (8u1e as reference) Cα distance vs. amino acid sequence.

*(b)* Overlay of chain B in 8u1e and the compared structure with residues colored by Cα distance.
